# Understanding the role of urban design in disease spreading

**DOI:** 10.1101/766667

**Authors:** Noel G. Brizuela, Néstor García-Chan, Humberto Gutiérrez Pulido, Gerardo Chowell

**Affiliations:** Scripps Institution of Oceanography, University of California, San Diego, La Jolla, California, USA; Department of Physics, Universidad de Guadalajara, Guadalajara, Jalisco, Mexico; Department of Mathematics, Universidad de Guadalajara, Guadalajara, Jalisco, Mexico; Department of Population Health Sciences, School of Public Health, Georgia State University, Atlanta, USA; Division of International Epidemiology and Population Studies, Fogarty International Center, National Institute of Health, Bethesda, Maryland, USA

## Abstract

Cities are complex systems whose characteristics impact the health of people who live in them. Nonetheless, urban determinants of health often vary within spatial scales smaller than the resolution of epidemiological datasets. Thus, as cities expand and their inequalities grow, the development of theoretical frameworks that explain health at the neighborhood level is becoming increasingly critical. To this end, we developed a methodology that uses census data to introduce urban geography as a leading-order predictor in the spread of influenza-like pathogens. Here, we demonstrate our framework using neighborhood-level census data for Guadalajara (GDL, Western Mexico). Our simulations were calibrated using weekly hospitalization data from the 2009 A/H1N1 influenza pandemic and show that daily mobility patterns drive neighborhood-level variations in the basic reproduction number *R*_0_, which in turn give rise to robust spatiotemporal patterns in the spread of disease. To generalize our results, we ran simulations in hypothetical cities with the same population, area, schools and businesses as GDL but different land use zoning. Our results demonstrate that the agglomeration of daily activities can largely influence the growth rate, size and timing of urban epidemics. Overall, these findings support the view that cities can be redesigned to limit the geographic scope of influenza-like outbreaks and provide a general mathematical framework to study the mechanisms by which local and remote health consequences result from characteristics of the physical environment.

**Author summary:** Environmental, social and economic factors give rise to health inequalities among the inhabitants of a city, prompting researchers to propose ’smart’ urban planning as a tool for public health. Here, we present a mathematical framework that relates the spatial distributions of schools and economic activities to the spatiotemporal spread of influenza-like outbreaks. First, we calibrated our model using city-wide data for Guadalajara (GDL, Western Mexico) and found that a person’s place of residence can largely influence their role and vulnerability during an epidemic. In particular, the higher contact rates of people living near major activity hubs can give rise to predictable patterns in the spread of disease. To test the universality of our findings, we ’redesigned’ GDL by redistributing houses, schools and businesses across the city and ran simulations in the resulting geographies. Our results suggest that, through its impact on the agglomeration of economic activities, urban planning may be optimized to inhibit epidemic growth. By predicting health inequalities at the neighborhood-level, our methodology may help design public health strategies that optimize resources and target those who are most vulnerable. Moreover, it provides a mathematical framework for the design and analysis of experiments in urban health research.

## Introduction

Empirical studies have identified inter-city variations in the timing, intensity and severity of influenza-like outbreaks [1–4]. Aiming to understand the mechanisms through which city characteristics yield such health consequences, epidemiologists have resourced to a variety of methods. Epidemiological data reveal compelling statistical correlations, but do not resolve intra-city variations in health that are driven by lifestyle inequalities at the neighborhood level [2,4–6]. In contrast, agent-based computational models use massive mobility datasets to recreate the behavior of individuals as they interact and spread infections [7–9,11–14]. While this approach allows for household-level analyses and its elevated complexity makes it suitable for targeted experiments [15], it is not the best tool for the development of general strategies in public health [16]. Furthermore, the information necessary to calibrate agent-based simulations is not openly available for most of the world’s cities. Thus, despite the fact that epidemics have the potential to be seeded anywhere, these intricate models have been overwhelmingly applied to populations in the developed world. Meanwhile, the fundamental mechanisms that drive health inequalities within metropolitan areas remain elusive.

Urban design determines the densities and relative locations of housing, jobs and services inside a city. Consequently, it influences the transportation choices of the population [17–19] and thus helps shape interaction networks through which diseases are spread. For example, the agglomeration of jobs and services drives large fractions of a city’s population to gather in small fractions of its area, increasing contact rates between the residents of distant neighborhoods [20,21]. In what follows, we quantify the agglomeration of urban mobility using two Gini coefficients (0 ≤ *G_origins_, G_destinations_* ≤ 1) that use neighborhood-level census data to measure spatial inequalities in the area-density of housing and activities throughout a city. The extreme values *G* = 0 represent homogeneous distributions of the population (trip origins) or its activities (trip destinations); inversely, *G* =1 indicates that all housing or activities are concentrated within a single location. In epidemiological terms, *G* = 1 produces homogeneous mixing conditions, which allow individuals to interact indiscriminately with all other members of the population. Under this scenario, disease spreading may be modeled using simple ordinary differential equations where the total population *N* = *S* +*I* + *R* is split into susceptible *S*, infected *I*, and recovered *R* groups. However, reality deviates from these conditions (*G* ≠ 1) and successful models must introduce heterogeneous contact networks to link the members of a population [22].

Attempting to understand the heterogeneous mixing patterns across subsets of the population, observational studies have tracked people’s interactions in day-to-day settings. It has since become clear that contact rates and the resulting risk of infection can vary widely with age, sex, employment status and other characteristics [23–26]. However, the complex demographics and social settings that exist in urban environments make it difficult to reliably sample and characterize the interaction patterns between all identifiable subgroups. Therefore, unexplained variations within sampled groups can cause statistical uncertainty intervals to be ≥50% the magnitude of reported mean contact rates [24–26]. Among the numerous likely sources of contact rate variaibility, crowding driven by urbanization and mobility patterns has been suggested to increase the growth-rate and size of infectious outbreaks by observational and modeling studies alike [4,14].

In this article, we develop a mathematical framework that introduces urban geography as the leading-order component of a susceptible-infected-recovered (SIR) epidemiological model. Our methodology uses spatially-resolved census data to infer transportation patterns in cities and is meant to bypass the need for large mobility datasets in urban health simulations. In our model, individuals are grouped by age and place of residence, and are subsequently represented through probability density functions that describe their likely travel habits. This approach sacrifices the hyperrealism of agent-based simulations to instead resolve the spatial patterns that arise when a heterogeneous set of metapopulations interact. Our goal is not to make an operational forecasting tool, but instead to formulate a scheme that helps understand the health consequences of the complex, spatially-dependent demographics of large cities.

As a first step, we characterize the role of urban design in this process and use the Guadalajara Metropolitan Area (GDL) as an example. City-wide hospitalization data from the 2009 Influenza A/H1N1 pandemic were used to calibrate disease parameters (S1 Fig) but cannot inform the validity of neighborhood-level patterns in our results. Lastly, we ran simulations in hypothetical cities with the same area, population density, number of schools and businesses as GDL. This allowed us to demonstrate that changes in the spatial distributions of housing, education and economic activities yield large variations in the size and early growth rate of epidemics across cities that would be deemed identical from a large-scale perspective.

## Modeling framework

### Inferred mobility patterns

High-resolution maps of residential density provide the background distribution of a city’s population, introducing a first layer of spatial dependence to the SIR transmission framework. Likewise, the locations and prominence of activity hubs determine common trip destinations were human interactions occur and diseases are spread. In our gravity model, trips originate at home and the probability density *P*(x, y′) that a person residing at x will visit y′ on a given day is the joint result of two factors: The distance Δ*r_xy′_* between both sites and the overall popularity of y′ as a destination. The effect of distance Δ*r_xy′_* on the likelihood of displacements between two points has been studied empirically [27], yielding the general form

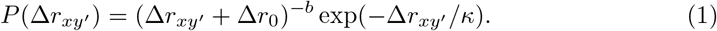

Simplifying daily mobility as radial displacements out of a place of residence x, *P*(Δ*r_xy′_*) yields a first estimate of the spatial distribution of individuals as they go through their daily routines. Moreover, the model parameters Δ*r*_0_, *b*, *κ* can be used to capture the effects of transportation infrastructure, as the quality of these services largely determines the willingness of individuals to travel long distances [28].

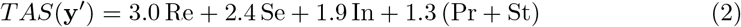

Trip Attraction Strength (TAS, Eq. 2) further refines our approximation of human mobility patterns in cities. TAS is an estimate of the daily number of visitors driven into a location y′ by education and economic activities [29]. Its value may be calculated using school enrollment data and the number of workers employed at y′ by different economic sectors, allowing different types of establishments to be weighed by the traffic they each induce. Our model incorporates TAS through Equation (2), where parameters Re, Se, In and Pr denote the number of jobs registered at y′ by retail, service, industrial and primary activity organizations respectively [29]. Similarly, St is the number of students enrolled at educational institutions inside the same area.

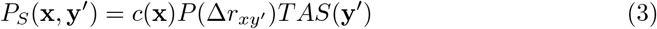

Equation (3) thus defines a probabilistic gravity model that represents mobility as the weighted influence areas of individuals who try to minimize their displacements but will travel farther given an economic incentive (higher TAS). Here, the normalization factor *c*(x) allows to adjust for the average number of daily trips made by residents of different neighborhoods. In what follows, we use the empirical values *κ* = 80 km, *b* = 1.75 found by González and collaborators [27] but prescribe Δ*r*_0_ = 5. This is a conservative choice, as it facilitates longer trips and thus homogenizes the transportation habits of the city’s inhabitants.

Before integrating these concepts into an epidemiological transmission model, we must distinguish between the behavior of susceptible and infected groups. We define the mobility of symptomatic groups via Equation (4), where 0 < α(x) < 1 is an isolation parameter whose spatial dependence allows the representation of neighborhood-targeted intervention strategies as well as social and economic factors that influence the adaptive behavior of sick individuals. The term *H*(x, y′) accounts for the added probability of visits to the nearest healthcare facility, as sick individuals will likely seek diagnosis and treatment.

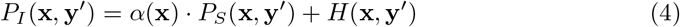

In summary, we use the gravity model in Equation 3 to infer mobility patterns in cities. *TAS* represents the number of people who visit a location on a given day and is calculated using employer and educational databases alike (Eq. 2, see Materials and Methods). Although the prominence of different destinations varies throughout the week and within a single day (for example restaurants and schools have strongly marked daily cycles), our simulations consider *TAS* to be fixed in time. Lastly, we incorporate all state-run healthcare facilities (henceforth hospitals) by assigning infective individuals to the nearest hospital and adding an 8% probability that they will visit it once during their infective period. An overview of all spatial components of our framework, as estimated for GDL (see Materials and Methods), is shown in Figure 1.

**Fig 1.**
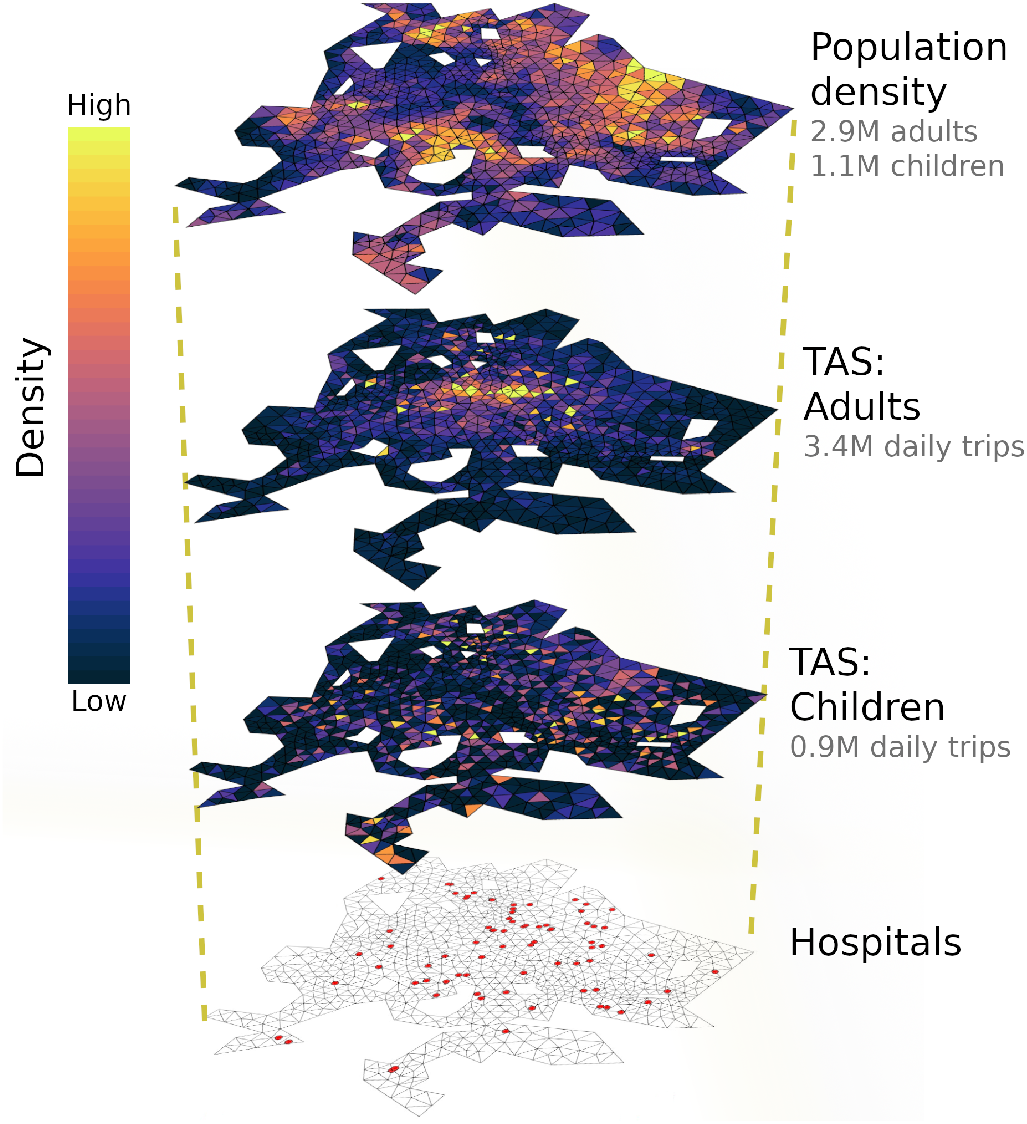
Layers of model input: population density and TAS were calculated for adults and children within a triangular grid representing GDL (Eq. 2, see Materials and Methods). While children are attracted to early-stage schools, the destinations of adult mobility are determined by the locations of retail stores, factories, office buildings, high schools and universities. Satellite image: Google, Landsat/Copernicus, Maxar Technologies.

## Metapopulation model

Let us define three identical domains 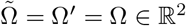 to represent the urban region under study. Members of the population *N* are then segregated under an SIR transmission scheme and distributed following population density functions *S*(x, *t*) + *I*(x, *t*) + *R*(x, *t*) = ρ(x) such that 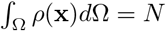. This approach links people to their place of residence (henceforth x ∈ Ω for susceptibles and 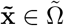 for infectives), allowing to formulate a Distributed Contacts SIR model (DC-SIR, Eqs. 5–7 adapted from [30,31]). Here, *β*(y′) is the probability of contagion given a susceptible-infective interaction that happens at y′ and *γ* is the recovery rate. The interaction kernel *k*(x, y′,*t*) yields the expected number of interactions between a member of *S*(x, *t*) and all infected individuals present at a trip destination y′ ∈ Ω′ and time *t* [30,31]. Thus, *k*(x,y′,*t*) introduces the heterogeneous contact networks through which diseases are spread.

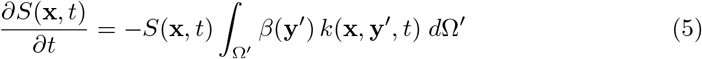

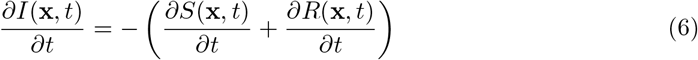

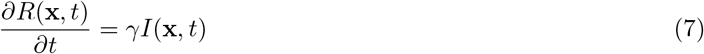

To define the interaction kernel *k*(x, y′, *t*), we use origin-destination density functions *P_S_*(x,y′,*t*),*P_I_*(x,y′,*t*) for members of *S*(x,*t*) and *I*(x,*t*) respectively. Next, we compute the total number of infected individuals expected to visit y′ at a given time, which is the integral from all trip origins 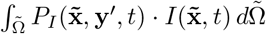. Taking the product with *P_S_* (x, y′,*t*) thus gives an expression for the expected number of SI contacts occurring at y′ for an individual who resides at x

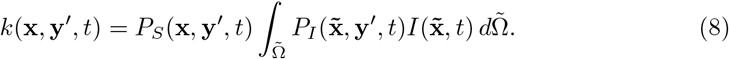

A similar procedure yields the reproductive number 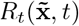, defined as the number of secondary cases caused by a single infected individual who lives at 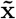 and contracted the disease at time *t*. Instead of calculating the number of SI interactions for a susceptible in transit, we now use the spatial distribution of susceptibles at a given time. This is given by the integration of susceptible mobility over all trip origins 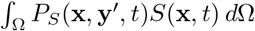. Multiplying this by the origin-destination density for infected mobility *P*_I_(x, y′,*t*) then yields the expected number of SI interactions occurring at y′ for a member of 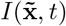

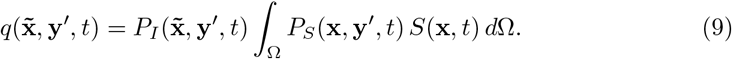

Assuming that all variables are slowly-varying over the first generation of disease transmission, we can approximate the basic reproduction number 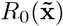 as

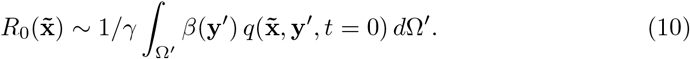

Notice that *k*(x, y′,*t*) and 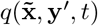 represent the vulnerability and disease spreading capacity of individuals during an infectious outbreak. Moreover, their spatial dependence implies that a person’s role during an epidemic is a function of their place of residence but determined by characteristics of the locations they visit. With this mathematical foundation, one could consider developing parameterizations of environmental conditions such as relative humidity (function of y′) and demographic factors that influence contact patterns (functions of x, 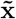).

Our DC-SIR simulations of GDL classify two subsets of the population by place of residence and age group. Thus, we defined SIR subgroups *S_j_*, *I_j_*, *R_j_* and mobility functions *P_Sj_*, *P_Ij_* to represent the people in each age group *j* = 1, 2 (adults and children). Consequently, the integrand over trip destinations Ω′ in Equation (5) was modified to account for interactions between a particular susceptible subgroup and infectives Ij of all ages, each of them with an age-specific transmission potential *β_j_*. Because our current focus is to explain the observed epidemiological impacts of urban geography and crowding [2,4,14], contact rates in our model are a linear function of visitor density. Although this assumption can fail to quantify face-to-face contacts within dense gatherings [32,33], the contact rate variations that emerge in our simulations of GDL are comparable to the statistical uncertainty of observational estimates [23, 25, 26] that have not offered a systematic explanation of this variability.

## Results

Our modeling framework was tested using data for GDL, where census and economic data [34–36] were processed to derive the daily mobility patterns of children (age ≤ 15) and adults (age > 15, see Materials and Methods). Lorenz curves in Figure 2 represent the different degrees of agglomeration driven by housing (*G_origins_* = 0.32) and daily activities (*G_destinations_* = 0.56). As is true for large cities [20,21], inferred trips were largely directed towards a few, hyper-affluent areas. In our model, these areas receive 25% of all daily trips but occupy less than 5% of the metropolis; meanwhile, housing places 25% of the population throughout 13.5% of the city’s most densely populated regions. The condition *G_destinations_* > *G_origins_* requires that some neighborhoods have a net loss of occupants during workdays, as their inhabitants leave and agglomerate around major activity hubs. This is true for 60% of all neighborhoods in GDL, which then act as net sources of mobility whose role is to supply more affluent destinations with visitors.

**Fig 2.**
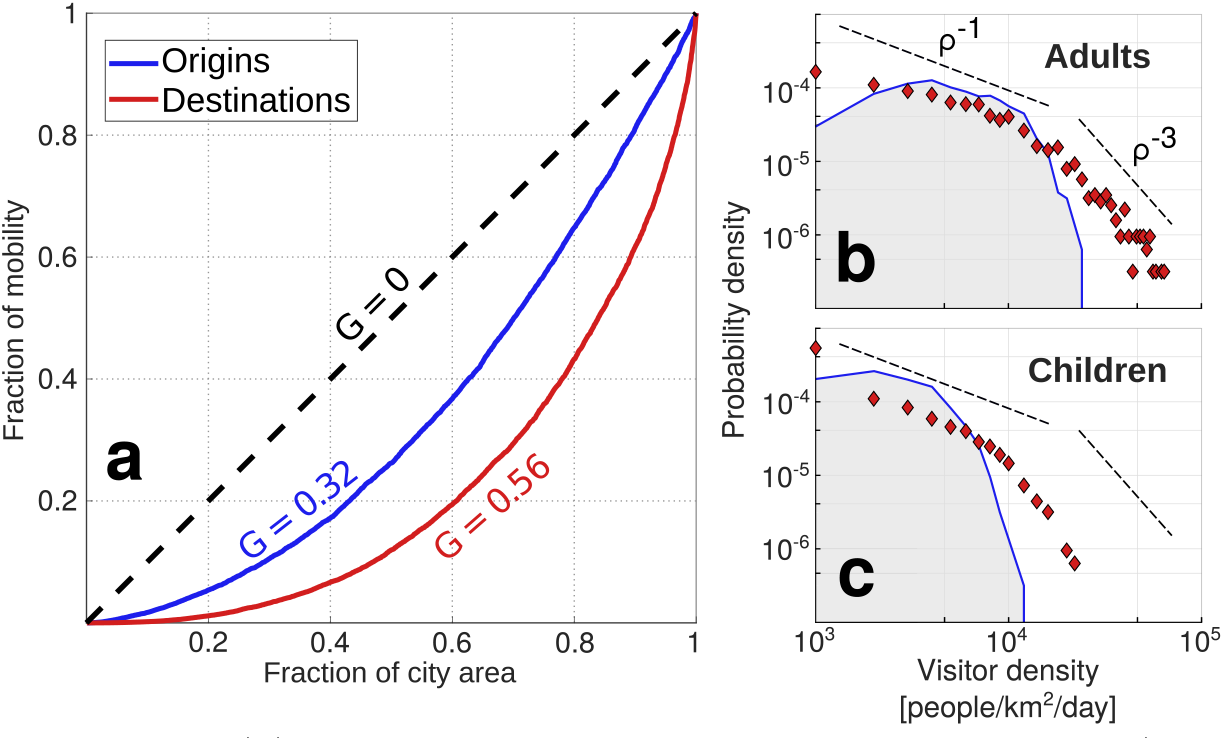
Lorenz curves (a) show the cumulative distributions of housing (trip origins, blue) and daily activities (trip destinations, red) throughout portions of the city area. The dashed line (*G* = 0) here represents urban design with spatially-homogeneous distributions of housing and daily activities. Probability density functions of visitor density (note the logarithmic axes) show that daily activities (red diamonds) drive greater crowding than housing (blue line) for both adults (b) and children (c).

Values of *β*(y′) were calibrated using city-wide hospitalization data from the 2009 A/H1N1 influenza pandemic (S1 Fig). In our model, this parameter yields a linear relationship for the probability of falling ill given the mean area density of infectives that one encounters throughout the day. We assumed a homogeneous transmission rate *β* = *β*(y′) and obtained that the daily risk of contagion increased by 1.1 ± 0.1% for every 1000 infective adults per *km*^2^ added to one’s surroundings. Similarly, the addition of 1000 infective children per *km*^2^ raised the probability of falling ill on a given day by 3.2 ± 0.3%. With β fixed everywhere, spatial patterns in the evolution of epidemics result entirely from the number and age of people who visit each one of the city’s neighborhoods (Eqs. 8, 9). Hence, spatial variations in our simulation results and in the basic reproductive number 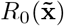 are a direct consequence of the city’s transportation network (Eq. 10), which is by definition a product of urban design and land-use patterns (Eqs. 1 – 3).

Spatial variations in the basic reproductive number 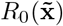 (Fig. 3) had a profound impact on the spatiotemporal evolution of outbreaks in our model. Firstly, the age and place of residence of patient zero (initial conditions) significantly influenced the early rate of epidemic growth. In fact, the *R*_0_ of patient zero can delay the peak of an epidemic by as much as 9 days (Fig. 4.a,b). Secondly, people with the highest 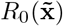 also had the highest probability of falling ill. In particular, the increased contact rates of downtown residents allowed them to spread pathogens 50% more efficiently than their suburban counterparts (Fig. 3), but also put them at a higher risk of contagion. Regardless of initial conditions, this particular situation gave rise to a spatial pattern in which influenza spread as waves of disease that emanate out of the city center (Fig. 4.c, S2 Fig, S1 Video). Similar patterns have been observed in agent-based simulations before [10] and may thus be a general characteristic of flu-like epidemics in urban populations.

**Fig 3.**
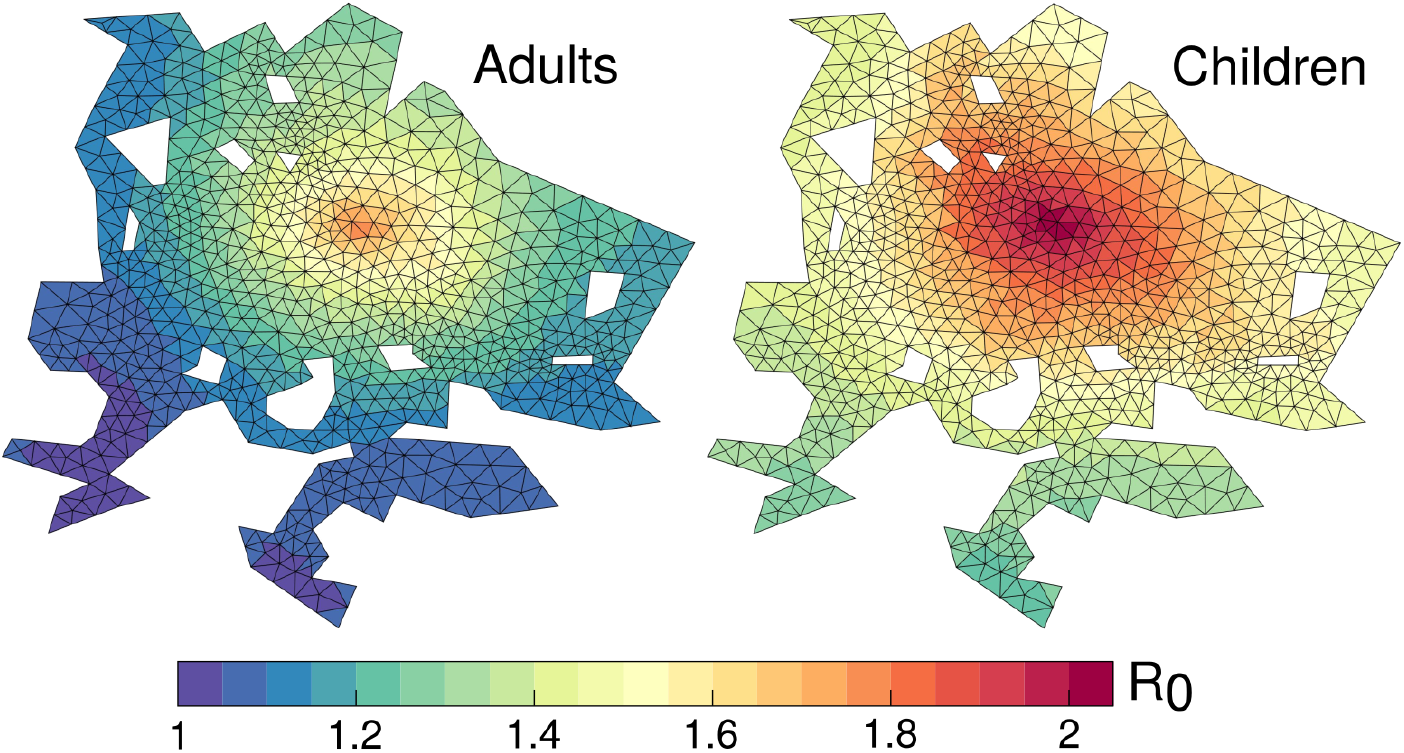
The basic reproductive number *R*_0_ (Eq. 10) varies at the neighborhood scale, showing that adults (left) and children (left) living near the city center play a disproportionate role in disease spreading. Population-averaged values of *R*_0_ were 1.27 for adults and 1.57 for children, and scale linearly with contact rates in our simulations (Eq. 10), meaning that the absolute range of contact rates for both age groups was roughly 50% of the mean value.

**Fig 4.**
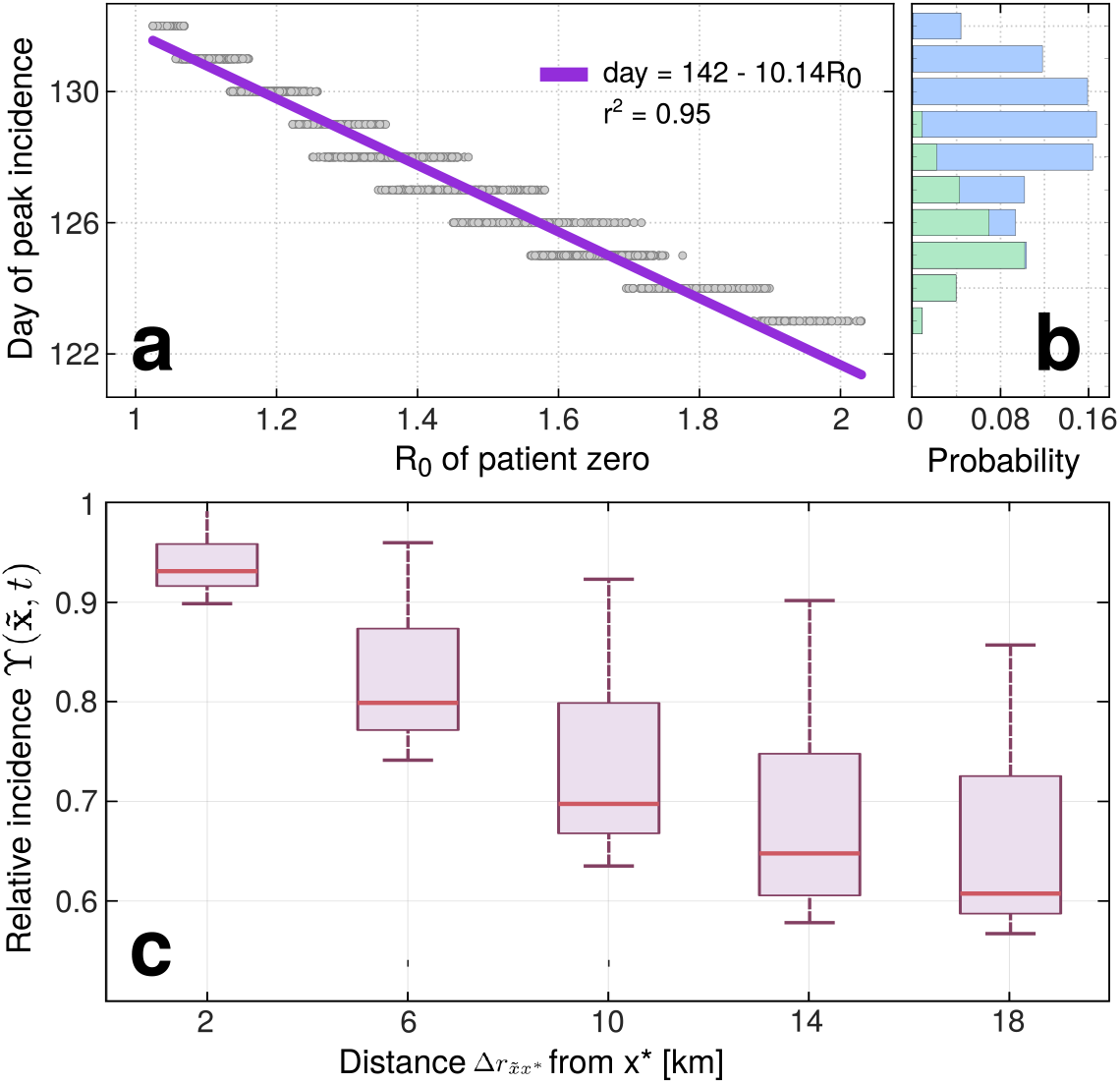
Over 3160 simulations with different initial conditions, panel a shows the time of peak incidence (when 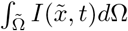 reaches a maximum) plotted against the *R*_0_ of patient zero. A histogram in panel b shows the probability that an outbreak peaks on a given day and the separate contributions of children (green) and adults (blue) to this variability. Boxplots in c show radial decrease in expected (1, 25, 50, 75 and 99th percentiles) relative incidence 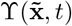 at a location 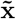 given its distance 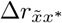 to the city’s largest retail hub x*.

To better appreciate the structure of wave-like patterns and inequalities in the spread of disease, we define the relative incidence 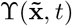 (Eq. 11) and use simulation results to inspect its statistical dependence on the distance 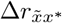 between neighborhoods 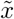 and the city’s largest retail hub x*, where TAS peaks.. Boxplots in Figure 4.c show that our analyses predict the incidence of influenza to be highest in downtown GDL and decreasing towards the suburbs. Time dependence represented in S2 Fig suggests a relationship of the form 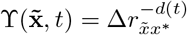. Here, *d*(*t*) is a non-negative function that decreases with time, as 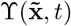 approaches unity everywhere towards the end of outbreaks.

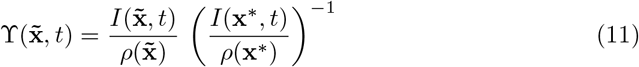

It is clear from Figures 3 and 4 that mobility hotspots play an important role in setting the growth rate and evolution of epidemics in our model. To gain more insight, we relocated businesses, schools and housing across the GDL numerical grid and ran simulations in the resulting geographies. This allowed us to compare outbreaks across cities with unique transportation networks but the same demographics, area, number of schools, businesses and daily trips as GDL. In one set of experiments we modified the Gini coefficients of mobility by relocating businesses, schools and housing across areas with low and high TAS or population density. As a result, we modulated the agglomeration of urban mobility but preserved the spatial structure of GDL. In another experiment, we redistributed housing, schools and businesses at random to evaluate whether the effects of agglomeration can be generalized to all cities despite their spatial structure. All simulations used the same values of *β* (previously calibrated for GDL) and an infective period 1/*γ* of 4 days. Results are summarized in Figure 5.

**Fig 5.**
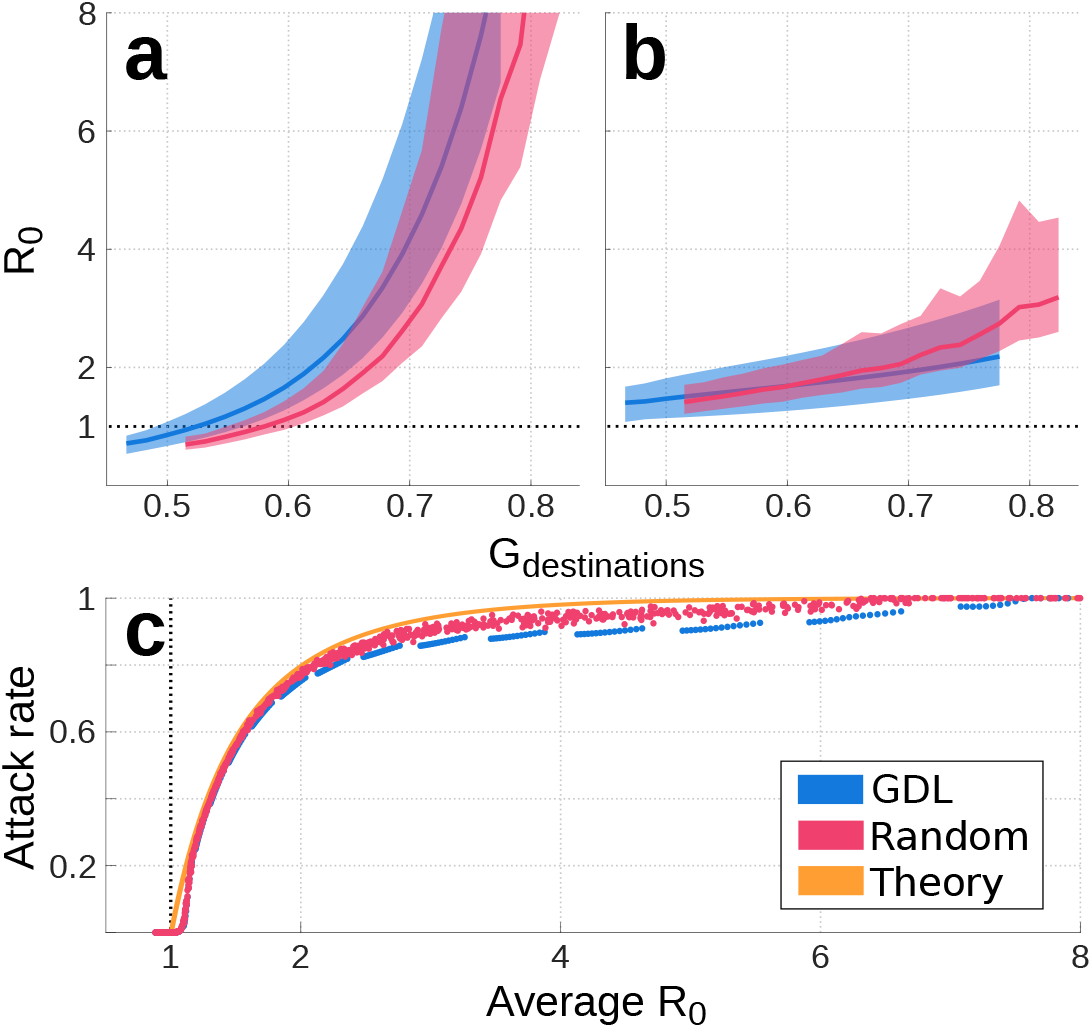
The population-averaged basic reproduction number *R*_0_ is plotted as thick lines for adults (a) and children (b). Color shading shows the range of values 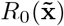 calculated with equation (10) in cities with the spatial structure of GDL (blue) and those with randomized urban design (pink). The relationship between population-averaged *R*_0_ and attack rate *z* is shown in panel c for cities with realistic (blue) and random (pink) spatial structures. The theoretical relation for the attack rate under homogeneous mixing conditions (Eq. 12) is shown in orange. Disagreements between these curves show the impacts of urban design on the growth rate and size of influenza-like outbreaks for cities with the same area, population and economic activities.

Our simulations suggest that, for all cities, the basic reproduction number *R*_0_ varies as a function of *G_destinations_* and is virtually independent of *G_origins_* (Fig. 5.a,b). This relationship is nonlinear for adults and linear for children, mainly because *G_destinations_* is set by commercial and economic superclusters that primarily attract adults. Furthermore, our results suggest that housing and activities in metropolitan areas with *G_destinations_* < 0.58 can be redistributed so that the mean basic reproduction number of adults remains under the threshold value *R*_0_ = 1. Namely, in this illustrative case where *β* is fixed everywhere and transmission dynamics are solely driven by urban design, cities can be redesigned to render their populations incapable of sustaining epidemics.

At the end of an outbreak, the attack rate *z* measures the fraction of the population that contracted the disease. Under homogeneous mixing conditions (*G_destinations_* = *G_origins_* = 1), *R*_0_ and *z* are linked by Equation (12) [37]. At all values of *R*_0_, this relationship predicts higher attack rates *z* than observed in our simulations (Fig. 5.c). Disagreements are largest for low values of *R*_0_, when fractions of the urban population have *R*_0_ < 1 and thus do not contribute to exponential growth. This highlights the importance of understanding population heterogeneity: because *R*_0_ and incidence may covary in space (Figs. 3, 4.c), population-averaged parameters may differ from the characteristic values obtained from public health reports, whose samples are biased towards subgroups with a higher incidence. Differences between the GDL and Random experiments in Figure 5 suggest that, at equal *R*_0_, realistic urban layouts (where activity areas are clustered near each other) may act to decrease the size of epidemics when compared to cases where housing and activities are distributed randomly. However, notice that a greater *G_destinations_* is required for randomly-designed cities to have the same values of *R*_0_ as GDL (Fig. 5.a).

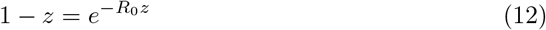

## Discussion

Our parameterization of contact heterogeneity in metropolitan areas produces smaller epidemics than homogeneous mixing models (Fig. 5) and predicts that, depending on where they live, people play unequal roles in disease spreading (Fig. 3). These inequalities relate the identity of patient zero to the timing of influenza-like outbreaks (Fig. 4.a,b) and can give rise to spatiotemporal patterns in the spread of disease (Fig. 4. c, S2 Fig, S1 Video). In fact, these processes mirror those of large scale transportation networks, whose heterogeneity leads to spatial inequalities in the size, timing and growth rate of epidemics [1,38,39]. As similar insight is used for the development of optimal intervention strategies in global-scale infectious outbreaks [40], results like those presented in Figure 3 may offer valuable information for public health officials who seek to optimize resources during infectious outbreaks. For example, health inequalities inferred here suggest that the efficiency of vaccination campaigns is likely to vary whether they target the inhabitants of the city center, its visitors, or people living in suburban areas (Figs. 3, 4). Similarly, the need for treatment and diagnosis is expected to differ across neighborhoods (Fig. 4.c).

Although our figures present the health impacts of urban geography as a function of place of residence, inequalities result primarily from the remote influence of people in other neighborhoods and the characteristics of places where interactions occur (Eqs. 5–10). The inclusion of non-residental processes is essential to the accurate representation of environmental health impacts [41,42] and is achieved here through interaction kernels *k*(x, y′,*t*) and 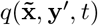. While we used a constant disease parameter, the representation of local and remote environmental conditions can be further refined by assigning full spatial dependence to transmission parameters so that 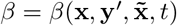. Generally speaking, our analytical framework is designed to map geographic information onto social space, establishing connections that are weighed by a risk of contagion *β*. This assigns realistic network characteristics to a spatially-explicit disease transmission model, which simplifies the interpretation of simulation results and may enable additional theoretical and statistical analyses [43,44].

Our results highlight the extent to which the assumption of homogeneous mixing in metropolitan areas can bias model results. Most notably, epidemic size in simulations that use the same transmission parameter *β* throughout hypothetical, seemingly identical cities, covers the entire range 0 < *z* ≤ 1 (Fig. 5.c). This suggests that macroscopic quantities such as population size and density hold little to no dynamical significance in the evolution of infectious outbreaks. Instead, the spatial organization of human mobility (quantified here using the Gini coefficient *G_destinations_*, Fig. 2) seems to control epidemic growth (Figs. 5.a,b) and is statistically correlated with population size [4,14]. Dalziel and collaborators explored this notion [14] by comparing agent-based model results across 48 Canadian cities, but were not able to isolate the effects of spatial heterogeneity from all other information embedded in massive mobility datasets that were used as model input.

Empirical relationships between *G_destinations_*, *R*_0_ and the attack rate z shown in Figure 5 suggest that, through its influence on transportation networks, urban design may be optimized to modify epidemic growth rates and thus reduce the probability of seeding large outbreaks. The existence of invasion thresholds under which populations cannot sustain epidemic growth is a general characteristic of metapopulation models [45]. While many authors have conjectured about the possibility of designing cities to minimize disease prevalence [4,14,46,47], we present the first dynamical argument to link these ideas and the invasion threshold proposition of Colizza and Vespignani [45]. More specifically, our results suggest that, by evenly distributing activity hubs throughout a city (instead of clustering them in the city center), city planners can segregate subsets of the population and potentially inhibit the rapid transmission of pathogens across distant neighborhoods (Fig. 5). Consistent with known statistical relations between the transmission potential of influenza and the spatial organization of human behavior [4,14], our framework explains a plausible mechanism behind health inequality across the neighborhoods of the worlds’ cities.

The mechanism through which urban design impacts epidemic growth is illustrated in Figure 6: when daily activities are centralized in a handful of hyper-affluent areas (Fig. 6.a), mixing patterns become more homogeneous [21] and thus favor interactions between residents of distant neighborhoods (Fig. 6.c). However, when activity hubs are distributed throughout the city (Fig. 6.b), economic opportunities become locally available to greater fractions of the population. Ultimately, this reduces the probability of interactions between residents of distant neighborhoods (Fig. 6.d) and can thus inhibit the spatial spread of influenza-like pathogens. This mechanism is evidenced by results in Figure 5. Positive relationships in Figure 5 demonstrate that the size and *R*_0_ of urban epidemics increase with the centralization of daily activities. Moreover, the results of simulations made using randomnized activity layouts (pink lines) show that when trip-attraction clusters are broken apart and their places of interest distributed randomly, *R*_0_ becomes significantly smaller even if *G_destinations_* remains constant (Fig. 5.a).

**Fig 6.**
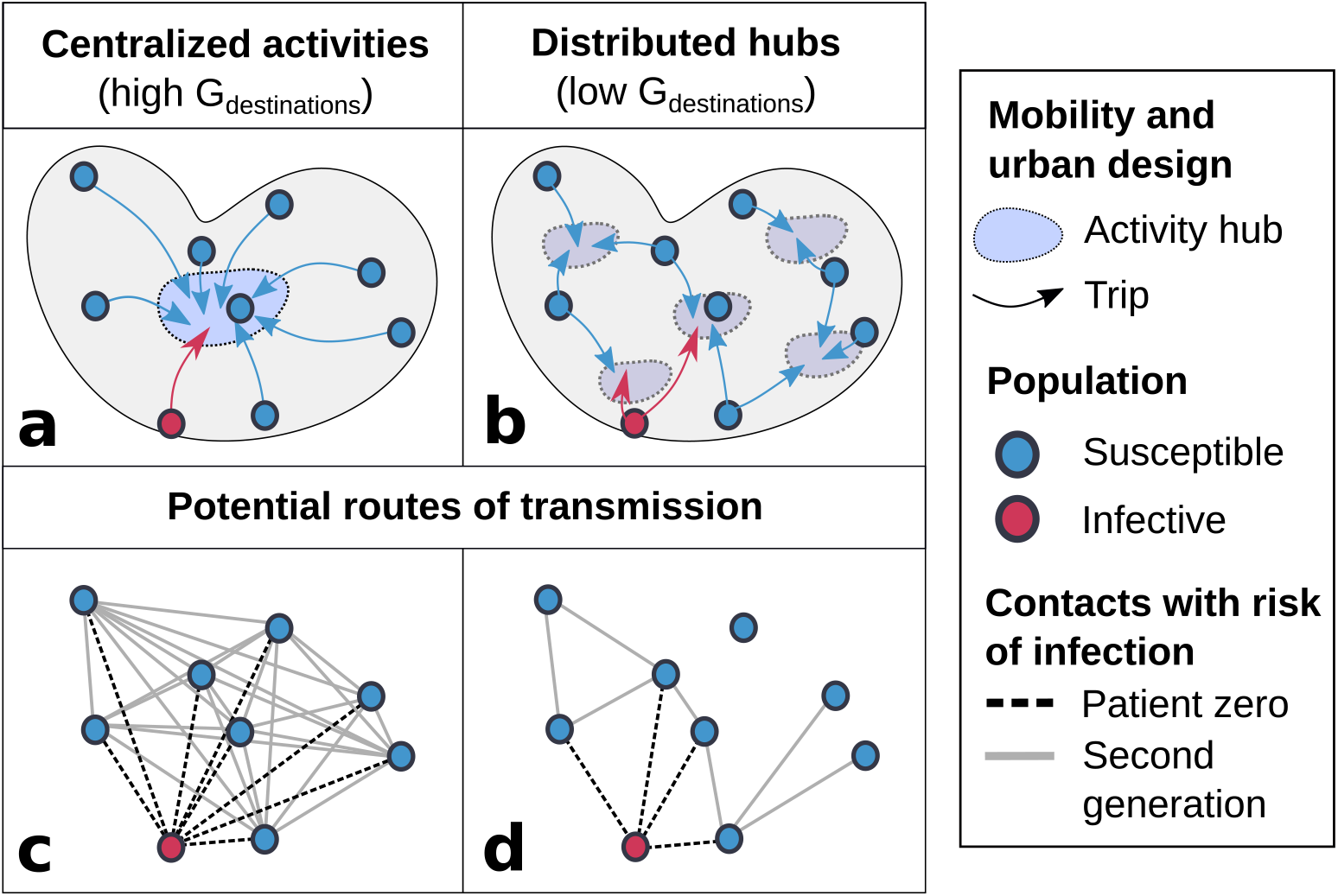
Schematic comparison of urban design scenarios. When activities are centralized in a single hub (left), people from all parts of the city gather in this place and can potentially interact [21]. Meanwhile, more even distributions of daily activities (right) segregate subsets of the population and can lower the growth rate of infectious outbreaks (Fig. 5).

Notice that children’s *R*_0_ was not affected by the breakup of activity hubs in Figure 5. b. This lack of effect may be due to the fact that children’s TAS is already scattered through GDL in a seemingly-random manner, while the TAS of adults is highly organized (Fig. 1). While the centralization of labor markets is thought to favor economic efficiency [19,48], our theoretical analyses suggest that this may have negative epidemiological consequences. In contrast, the decentralization of economic activities may help slow the propagation of influenza-like outbreaks (Figs. 5, 6), reduce pollution and improve overall welfare [49].

Agent-based models are the standard method to run realistic simulations of disease spreading in cities [7–10, 12–14]. These intricate models have allowed for the evaluation of intervention strategies [12] and health inequalities [8,15], but require input from massive mobility datasets that are not openly available for most of the world’s cities. Consequently, US [50] and European cities are over-represented by agent-based methods, which remain inaccessible for many research groups and public health organizations. Likewise, large-scale epidemiological models often assume homogeneous mixing conditions in cities and neglect all processes that occur at the neighborhood level [1,4,11,51]. When compared to agent-based model output, this simplification can lead to overestimate the size of epidemics and miscalculate their timing, and can thus introduce considerable bias [11]. Although far simpler, our parameterization of small-scale heterogeneity leads to the same conclusions (Figs. 4, 5). Thus, we believe that the mathematical framework presented here is an adequate alternative to introduce neighborhood-level processes in large scale epidemiological simulations.

Because it uses standard census and economic data to infer daily mobility patterns (see Materials and Methods), our method enables researchers to investigate transmission dynamics in virtually any city. Unfortunately, mobility parameters used here (Eqs. 1–3) were not calibrated with real origin-destination data (which exist for GDL during the 2009 A/H1N1 pandemic [13] but are not openly available) and may thus be inaccurate. Nonetheless, our analyses rely on the patterns that arise from fundamental processes driven by the agglomeration of economic activities and highlight small-scale inequalities that are not resolved by most epidemiological datasets. Agent-based simulations of influenza outbreaks in the city of Buffalo [10] show spatiotemporal patterns (their Figure 9) that are very similar to those presented here (S1 Video, Fig. 3) and thus suggest that urban design may play a leading-order role in setting the dynamics of influenza-like outbreaks in cities throughout the world.

In reality, spatial inequalities in health result from the complex interactions of social, environmental and biological processes among which urban design is only one. Even though this article focuses on the underlying effects of urban design on mixing heterogeneity and epidemic growth, our mathematical framework can incorporate additional layers of complexity that exist in reality. Among other factors, realistic representations of asymptomatic periods, reactive behavior and the use of time-varying mobility matrices could enrich future analyses. Nonetheless, the development of theoretical frameworks that seek to understand these complex interactions is crucial to help design new policies and research that aim to improve health in cities [46,47,52]. Ultimately, this study shows that systematic spatial variations in the mobility of infective individuals can drive robust patterns in the spread of disease and thus give rise to health inequalities at the neighborhood level (Figs. 3, 4, S1 Video).

## Materials and methods

### Inclusion of census data

We solved equations (5)–(7) in finite differences by mapping all their variables onto a 1580-element triangular mesh representing GDL ([53], Fig. 1). Neighborhood-level data from the 2010 census [35] were used to calculate the total adult (age > 15) and infantile (age ≤ 15) populations at each grid element. *TAS*(y′) was estimated for adults and children from two publicly available datasets: The 2015 National Statistical Directory of Economical Units (DENUE) lists all registered employers in Mexico along with their sector, number of workers and location [34]. Similarly, the National System for School Information (SNIE) locates all of the country’s schools and universities and lists their enrollment at each educational stage [36]. DENUE employment data were combined with SNIE enrollment at the high school and university levels to calculate TAS for the adult population via equation 2. On the other hand, TAS for infantile mobility was inferred using SNIE enrollment data for educational stages up to the middle school level, roughly corresponding with the age threshold between our age groups. Next, time-constant origin-destination matrices were obtained for all grid elements using equations (3) and (4). Resulting maps of population density and TAS for adults and children are shown in Figure 1.

Mobility parameters were established a priori as Δ*r*_0_ = 5 km, *b* = 1.75, *κ* = 80 km in equation 1. Sensibility analyses showed that variations in these values primarily impact *G_destinations_*, whose consequences are shown in Figure 5, but do not change the fundamental phenomena described in this study. Similarly, *c*(x) in equation 3 was chosen to yield an average of 1.18 and 0.85 daily trips per person for adults and children respectively. The mobility of infected individuals was computed via equation 4 with *α* = 0.8 and an 8% probability of visiting the hospital once in an infective period.

### Numerical solutions

Time-varying solutions to equations (5)–(7) were obtained using a forward finite-difference scheme that considered piece-wise constant functions defined over the triangular mesh shown in Figures 1 and 3. Continuous input variables were transformed to be constant at all grid elements, which are henceforth noted as Ω_*j*_ ⊂ Ω, and whose area is 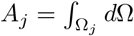. For instance, the number of susceptibles in Ω_*j*_ at time *t_k_*, was calculated as 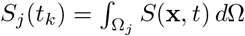. Origin-destination matrices were defined using the corresponding origin-destination density functions (Eq. 3) as

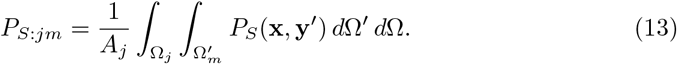

Similarly, *β*(y′) can be mapped onto the grid as

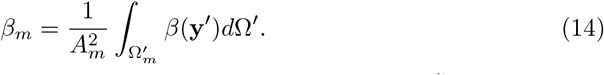

With all variables defined within the numerical grid, integrals over 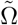 and Ω′ in equation (5) were replaced by sums over infective origins 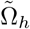 and trip destinations 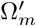. Thus, the temporal evolution in the number of susceptibles at Ω_*j*_ over one timestep Δ*t* was computed as

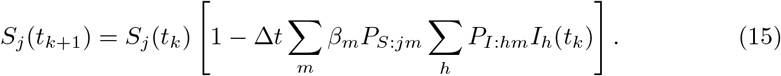

### Data and code availability

MATLAB software and data used to perform the simulations described in this article are available at https://github.com/inciente/Urban-Epidemiology/tree/master/RUDDS/. A Python Jupyter Notebook that describes our methods is also included.

## Supporting information

**S1 Fig. Comparison of data and model output**. The city-wide number of weekly hospitalizations in GDL during the 2009 AH1N1 influenza pandemic (blue) is compared to simulation results under various initial conditions (gray). Because our model does not feature interventions nor a time-dependent behavioral response to outbreak intensity, values of *β* were calibrated using only 9 weeks of data at the onset of the third and largest wave in the AH1N1 epidemic (pink shading).

**S2 Fig. Radial dependence of relative incidence**. Expected evolution of relative incidence at different times since the onset of 1580 outbreak simulations, each run under a different initial condition. Color shading shows probability density functions for different values of x*. At any time, the proportion of people who are sick is maximum near the city center where TAS (Eq. 2) peaks, and decreases radially towards the suburbs. As epidemics die out and the pool of susceptibles available to spread infection is depleted, relative incidence approaches unity everywhere, weakening the health contrasts that exist between residents of the downtown and suburban areas.

**S1 Video. Spatiotemporal evolution of an outbreak**. Model output shows the time-dependent fraction of people who are sick at each location in our numerical grid. The outbreak shown in this video was seeded by adding 15 infective adults to a grid element in the suburbs.

## Acknowledgments

The authors wish to acknowledge Haley McInnis, Thomas Gorin and Tarik Benmarhnia for their valuable feedback and suggestions. NB, NGC and HGP were supported by Consejo Nacional de Ciencia y Tecnología.

## Notes

#### Summary of Updates

We improved our discussion of existing literature on contact rates and inequalities in agent-based models. Figures 1-4 were modified to improve clarity and Figure 6 is an all-new schematic that summarizes the mechanisms responsible for our results.

https://github.com/inciente/Urban-Epidemiology/tree/master/RUDDS

