## Supplementary figures and images for "Understanding the role of urban design in disease spreading"

### S1 Fig

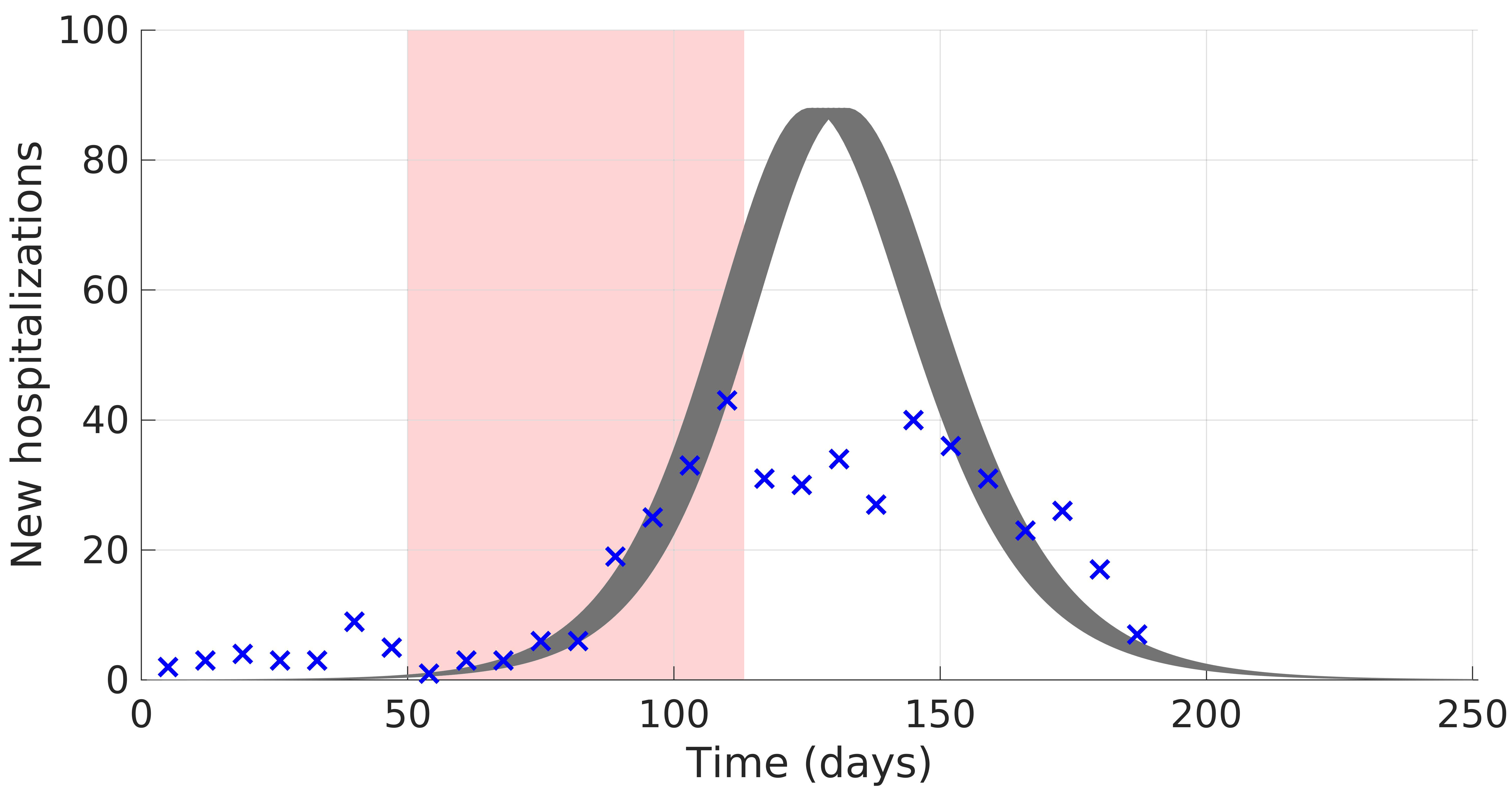

### S1 Video

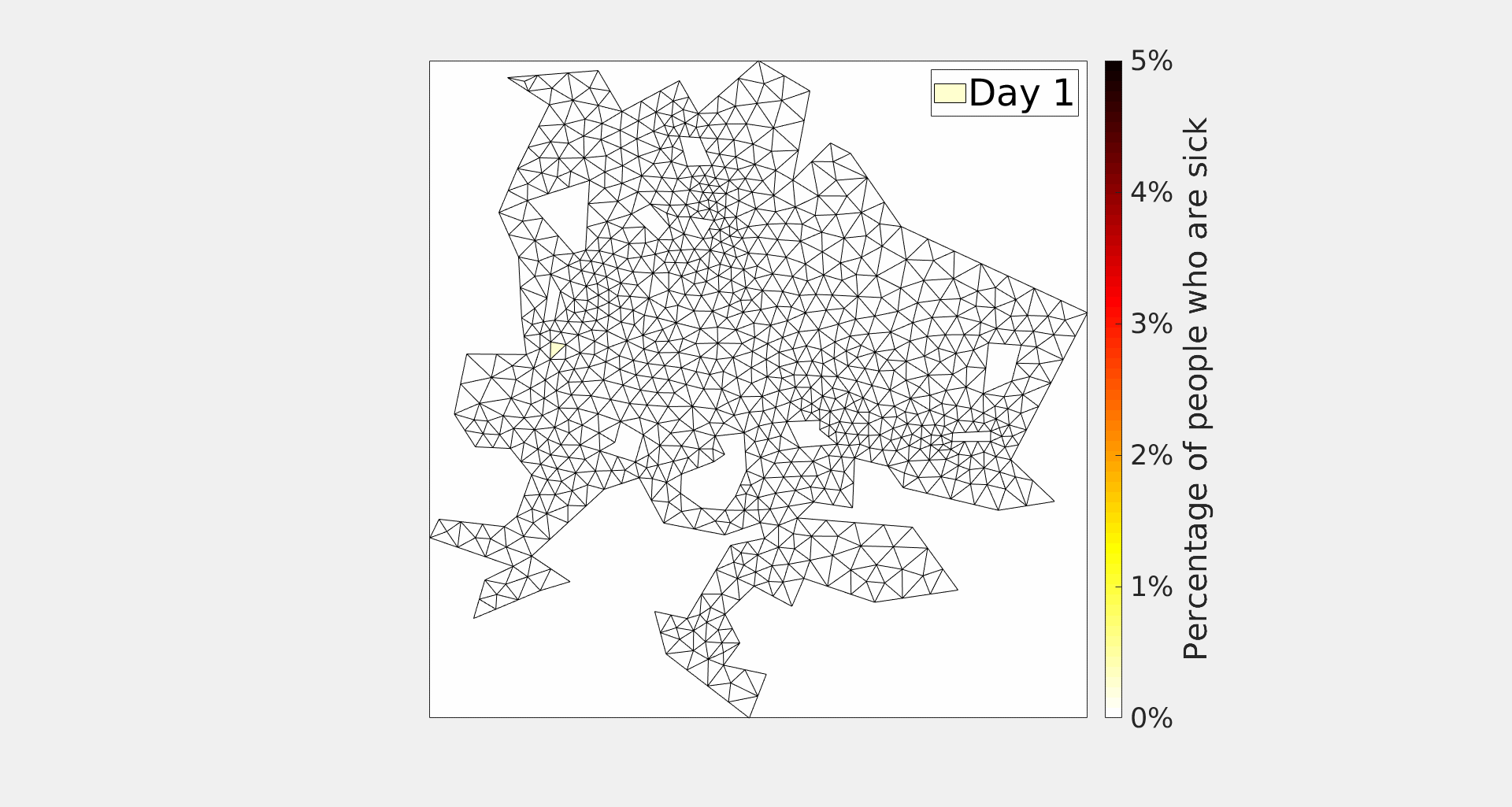

### S2 Fig

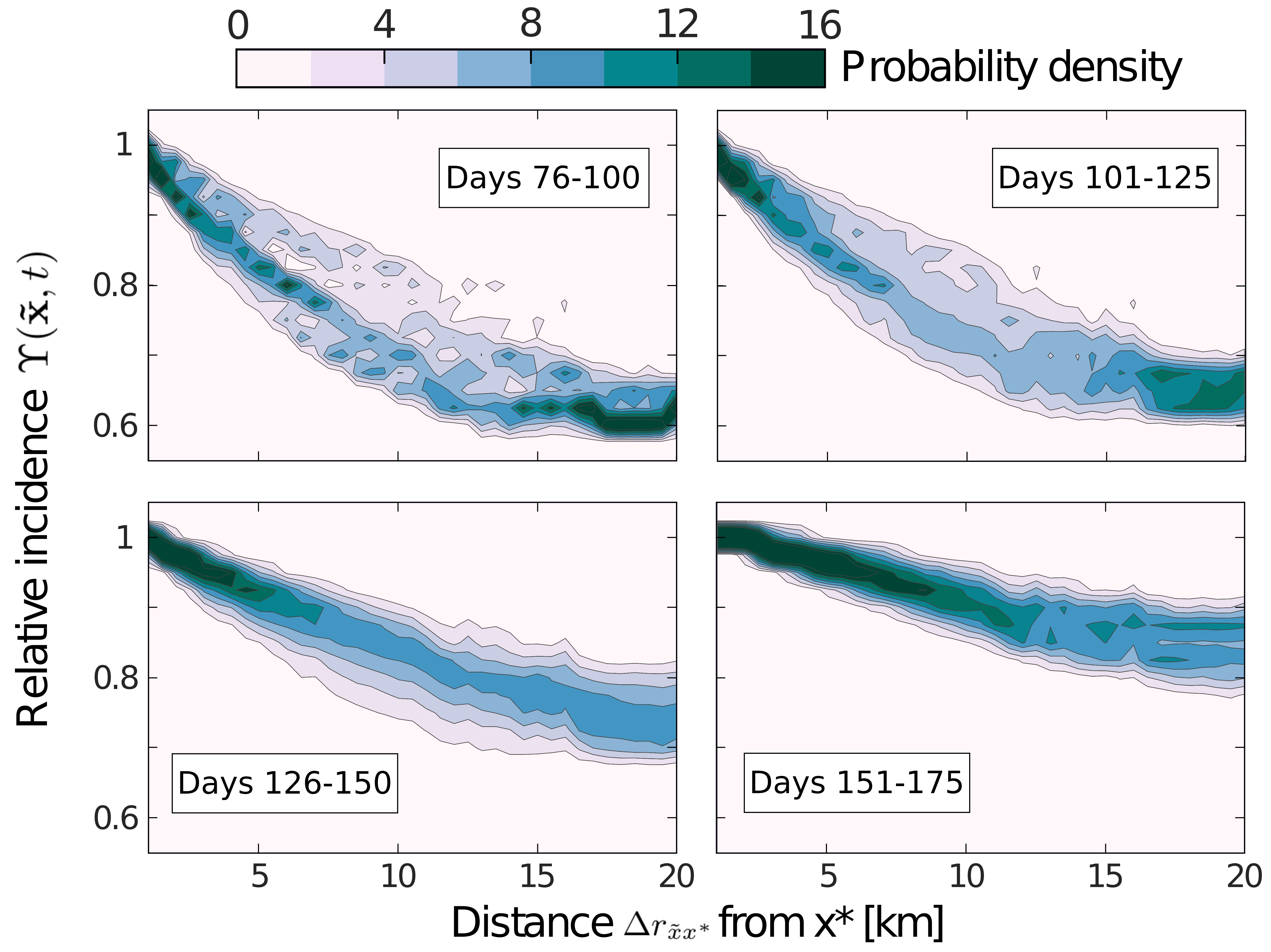
